# Functional Activity and Binding Specificity of small Ankyrin Repeat Proteins called Ankyrons against SARS-CoV-2 variants

**DOI:** 10.1101/2024.10.11.617752

**Authors:** Yun-Jong Park, Wojciech Jankowski, Nicholas C Hurst, Jeremy W Fry, Nikolai F Schwabe, Linda C C Tan, Zuben E Sauna

## Abstract

Effective management of COVID-19 requires clinical tools to treat the disease in addition to preventive vaccines. Several recombinant mAbs and their cocktails have been developed to treat COVID-19 but these have limitations. Here, we evaluate small ankyrin repeat proteins called Ankyrons™ that were generated to bind with high affinity to the SARS-CoV-2 virus. Ankyrons are ankyrin repeat proteins comprised of repetitions a structural module. Each module consists of a β-turn followed by two antiparallel α-helices. The Ankyrons™ are directly selected *in vitro* from a highly diverse library of around a trillion clones in ribosome display and like antibodies can bind with high affinity to almost any target. We assessed Ankyrons that were generated against the wild-type SARS-CoV-2 and the Delta and Omicron variants in a binding assay. We determined that all Ankyrons were specific in that they did not bind to MERS, a related coronavirus. While all Ankyrons bound with high affinity to the variant they were generated against, some also showed cross-reactivity to all three SARS-CoV-2 variants. Binding assays are useful for screening analytes but do not provide information about clinical effectiveness. Therefore, we used a pseudovirus-based neutralization assay to show that five of the Ankyrons evaluated neutralized all three strains of SARS-CoV-2. We have provided a workflow for the evaluation of novel Ankyrons against a viral target. This suggests that Ankyrons could be useful for rapidly developing new research tools for studying other emerging infectious diseases rapidly with the optional further potential for developing Ankyrons into diagnostic and even therapeutic applications.

**Author Summary:** Yun-Jong Park: Investigation, Methodology, Format Analysis, Writing – original draft.

Wojciech Jankowski: Methodology, Format Analysis, and Review.

Nicholas C Hurst, Jeremy W Fry, Nikolai F Schwabe and Linda C C Tan: Resources, Review & Editing.

Zuben E Sauna: Conceptualization, Supervision, Writing – original draft and Final editing.

## Introduction

The impact of severe acute respiratory syndrome coronavirus 2 (SARS-CoV-2), the virus responsible for coronavirus disease 2019 (COVID-19), on global health and the economy has been profound ^1^. The pathophysiology of COVID-19 poses unprecedented challenges, eliciting a robust immune response that can be exacerbated by a phenomenon known as “cytokine storm syndrome” ^2^. Additionally, SARS-CoV-2 infections can induce thrombogenesis which can result in multi-organ injury and can be fatal ^3^. Various drug classes, including antivirals, anti-SARS-CoV-2 antibody agents [monoclonal antibodies (mAbs), convalescent plasma, and immunoglobulins], anti-inflammatory drugs, immunomodulators, and anticoagulant drugs, have been proposed for the treatment and prevention of COVID-19 ^4, 5^. Biological molecules, such as mAbs, that block the SARS-CoV-2 spike protein from binding to the angiotensin-converting enzyme 2 (ACE2) receptor are clinically effective early in the disease ^6^. However, the safety concerns of biological therapy relate to immunogenicity [development of anti-drug antibodies (ADAs)] and possible immune potentiation effects ^7-10^.

An additional challenge in treating COVID-19 patients is that the SARS-CoV-2 spike protein, which is the primary target of mAb and vaccines, is prone to mutations. Thus, new virus variants periodically emerge that are not responsive to existing therapeutic antibodies and vaccines.^11^ To address this challenge, broadly neutralizing antibodies targeting the relatively conserved S2 subunit and its epitopes have been investigated ^12-14^. An approach that used a cocktail of two antibodies while initially successful in neutralizing SARS-CoV-2, was ineffective against new strains that emerged and the authorization for use was revoked.^15-17^ These experiences with conventional approaches indicate that there is a need to develop alternative therapeutic modalities that could neutralize diverse strains of SARS-CoV-2.

Proteins known as ankyrin repeat proteins are constructed with closely packed repetitions of approximately 33 amino acid residues^18^. Each repetition comprises a structural module that includes a β-turn followed by two antiparallel α-helices. A single ankyrin repeat protein may contain as many as 29 consecutive repetitions. However, four to six ankyrin repeats are more common and result in a right-handed solenoid structure characterized by a continuous hydrophobic core and an extensive, solvent-exposed surface^19, 20^. As a result, the binding surface takes on a groove-like configuration^21^. The ankyrin repeats used in the study described here are commercially produced and termed Ankyrons™.

Ankyrons™ are promising, non-immunoglobulin-based reagents that can be generated with immense diversity (>10^12^) and referred to as the Teralibary™ to bind almost any molecular target with high affinity. An Ankyron Teralibrary consists of fixed and variable positions. Fixed positions are important for structure, while each repeat module has a number of variable positions that are non-conserved, on the surface, and can potentially interact with the target.

Ankyrons are structurally different from antibodies with a single chain of linked ankyrin repeat binding domains. This makes Ankyrons suitable for creating multi-functional target binding reagents. Ankyrons also do not have many of the limitations of mAbs. The large size and complex structure of mAbs require advanced eukaryotic machinery for production, leading to high production costs. In contrast Ankyrons are as stable as fully formed IgG Antibodies while having only a single protein domain of around 15kD in size without relying on any disulfide bonding. The size of mAbs also presents challenges in reaching molecular targets that are not easily accessible. Moreover, the Fc region of mAbs interacts with multiple Fc receptors and can negatively regulate effector functions, impacting overall therapeutic efficiency. Unlike antibodies, Ankyrons have no known interactions with other receptors that modulate the immune system. However, like antibodies, Ankyrons are highly diverse, allowing the generation of a library with over a trillion clones. This library typically allows the direct selection of high affinity and highly specific binders against a target of choice in *in vitro* ribosome display without requiring further optimization such as molecular evolution. This approach is faster than other methods like phage display, which typically requires multiple rounds of screening and affinity maturation steps. Further, the diversity of library members that can effectively be displayed is only constrained by the number of ribosomes in the reaction which allows for a much greater diversity of library members to be used effectively compared to phage display. Ankyrons are about a tenth of the size (15kDa) of antibodies and their diverse encoding Teralibrary can be generated synthetically. Being selected directly *in vitro* from this synthetic Teralibrary, Ankyrons are not constrained by immunological tolerance and can be generated against all proteins, including self-proteins and proteins that are phylogenetically highly conserved. Additionally, there is no need to consider isotypes, as Ankyrons are small, single-domain proteins with high stability and heat resistance. Their stand-alone single domain functionality opens a vast spectrum of rapid engineering options for Ankyrons, far exceeding the scope available for antibodies, to deliver, e.g. bi-/multi-specific/-valent versions, and extended access to functionalization through chemical biology and orthogonal chemistries.

In the context of using DARPins, which, like Ankyrons, are also based on an ankyrin repeat platform, successful phase II results have been reported for ensovibep, a DARPin antiviral therapeutic for COVID-19. Ensovibep consists of five sequential, covalently linked DARPin domains, three of which have slightly different binding sequences (paratopes) to recognize overlapping epitopes of the receptor-binding domain (RBD) of the SARS-CoV-2 spike protein trimer.^22^ The combined binding of these three domains, each to a separate RBD, prevents the RBD from binding to the angiotensin-converting enzyme 2 receptor on host cells. The other two DARPin domains bind to human serum albumin to confer extended systemic half-life in vivo.^23, 24^

In our study, we utilize binding assays [using biolayer interferometry (BLI)] neutralization assays to evaluate the efficacy of Ankyrons, against both wild-type (WT) SARS-CoV-2 and various variants including Delta (B.1.617.2) and Omicron (BA.1). Our approach aims to understand the specificity of binding and the ability to prevent virus invasion in HEK293 cells that overexpress ACE2/TMPRESS. Unlike Ensovibep, which targets receptor binding on each RBD of the spike trimer using three domains, Ankyrons focus on a single domain, providing robust protection against a range of relevant spike mutants. In a viral passaging experiment, we demonstrate that the protective efficacy of single-domain Ankyrons against the development of viral escape mutants is comparable to that of a clinically evaluated mAb. Our findings suggest a potential new therapeutic approach that combines the strengths of mAb-based therapy while addressing the limitations.

## Results

### Assessment of Ankyrons against trimeric spike-RBD of SARS-CoV-2

Ankyrons specific to the following variants of the spike protein of SARS-CoV-2 were assessed: wild type (8 Ankyrons), delta (6 Ankyrons), and omicron (9 Ankyrons). We determined binding parameters using the BLI assay for all Ankyrons-proteins combinations. The FcγR protein was used as a negative control and the MERS trimeric spike protein was used to demonstrate that the Ankyrons are specific to the SARS-CoV-2 spike protein (Supplementary Fig. 1). In investigating protein-ligand interactions using BLI, it is crucial to effectively minimize nonspecific binding between Ankyrons and the biosensors, as well as between Ankyrons and spike proteins/control proteins. To do this, the spike protein was titrated (concentration range 0-50nM) on the biosensors and the concentration at which the slow initial binding is observed that does not saturate the biosensor was selected as the optimal loading concentration (Supplementary Fig. 2 and Supplementary Table 1). Several binding parameters are obtained with BLI, and binding response and KD have been used previously to measure ligand-analyte interactions ^25, 26^.

**Table 1.** Summary of Biolayer interferometry and Neutralization analysis for binding and inhibitory responses of SARS-CoV-2 Spike variants to Ankyrons. KD, equilibrium affinity constant; M, mol/L; >600nM of PsVNA50 (IC50) was not available due to poor binding with inaccurate curve fitting; a mean KD of <1E-12 M was not calculated for variants because of strong binding between analytes and ligands.

|  | WT | Delta | Omloron | WT | Delta | Omloron | WT | Delta | Omloron |
| --- | --- | --- | --- | --- | --- | --- | --- | --- | --- |
| WT-1 | 3.88 | 10.13 | 30.30 | 0.7041 | 0.0080 | 0.0105 | <1.0E-12 | 4.88E-08 | <1.0E-12 |
| WT-2 | 5.82 | 63.05 | 370.23 | 0.8243 | 0.0173 | 0.0118 | <1.0E-12 | 3.76E-08 | <1.0E-12 |
| WT-3 | 0.81 | 32.03 | 253.55 | 0.7005 | 0.0005 | 0.002 | <1.0E-12 | 5.87E+11 | <1.0E-12 |
| WT-4 | 0.88 | 31.54 | 324.15 | 0.81175 | 0.1194 | -0.0147 | 3.35E-11 | 2.21E-08 | <1.0E-12 |
| WT-5 | 18.12 | 66.01 | 800.00 | 0.81895 | 0.0028 | 0.0005 | <1.0E-12 | 3.26E-00 | <1.0E-12 |
| WT-6 | 1.48 | 187.80 | 138.03 | 0.4638 | 0.0028 | 0 | <1.0E-12 | <1.0E-12 | <1.0E-12 |
| WT-7 | 1.88 | 1.11 | 0.91 | 0.8033 | 1.381 | 1.121 | 1.13E-09 | 0.00E-10 | 1.94E-10 |
| WT-8 | 0.88 | 0.80 | 2.27 | 0.8838 | 1.4401 | 0.5257 | 1.32E-09 | 8.70E-10 | 1.24E-09 |
| DT-1 | 0.80 | 0.30 | 2.51 | 1.0428 | 1.7384 | 1.0722 | 1.07E-09 | 7.11E-10 | 8.08E-10 |
| DT-2 | 24.18 | 3.18 | 213.58 | -0.0282 | 1.8888 | 0.0158 | 4.13E+20 | 1.3E-09 | <1.0E-12 |
| DT-3 | 3.38 | 0.88 | 5.10 | 0.8828 | 1.31845 | 0.4348 | 1.33E-08 | 1.08E-08 | 4.84E-08 |
| DT-4 | 351.12 | 0.87 | 118.37 | 0.1862 | 1.82885 | 0.1189 | 8.97E-10 | 4.86E-10 | 7.33E-08 |
| DT-5 | 314.07 | 1.47 | 800.00 | 0.0457 | 1.82285 | 0.0384 | 1.89E-09 | 1.28E-08 | 3.07E-09 |
| DT-6 | 800.00 | 1.87 | 800.00 | -0.0021 | 0.9847 | 0.0002 | 1.18E-05 | 2.71E-09 | 1.06E-08 |
| Ome-1 | 800.00 | 124.44 | 1.96 | -0.0033 | 0.0035 | 1.20455 | 4.91E+09 | 5.30E-05 | 1E-09 |
| Ome-2 | 396.51 | 800.00 | 0.83 | -0.0167 | -0.0043 | 0.9101 | 2.44E+11 | 2.47E+19 | 3.43E-09 |
| Ome-3 | 6.17 | 184.63 | 0.78 | 0.2804 | 0.362 | 0.95035 | 1.88E-09 | 1.61E-08 | 5.77E-10 |
| Ome-4 | 800.00 | 800.00 | 0.95 | -0.0083 | -0.0031 | 0.59645 | 2.36E+15 | 1.23E-07 | 4.44E-09 |
| Ome-5 | 385.85 | 130.11 | 0.73 | 0.0008 | 0.0015 | 0.3 | 1.61E-05 | 4.84E+15 | 1.08E-07 |
| Ome-6 | 800.00 | 62.00 | 1.58 | 0.0136 | 0.0122 | 0.1037 | 3.42E+00 | 1.03E+18 | <1.0E-12 |
| Ome-7 | 800.00 | 481.48 | 0.58 | -0.0129 | -0.0089 | 0.5708 | 1.08E+11 | 1.72E+14 | 8.25E-09 |
| Ome-8 | 556.48 | 588.25 | 1.88 | -0.0125 | -0.0038 | 0.7884 | 2.31E+19 | 7.78E-08 | 1.92E-09 |
| Ome-9 | 5.52 | 2.58 | 1.31 | 0.4014 | 0.4813 | 0.2483 | 9.63E-10 | 9.76E-10 | 1.22E-07 |

The binding response of Ankyrons generated against wild type, Delta, and Omicron spike proteins are shown in Supplementary Fig. 3. All Ankyrons generated against the wild type, have binding scores for wild type spike protein that are significantly higher than those for the negative control (FcγR protein). However, Ankyrons WT7 and WT8 cross-react with the Delta and Omicron spike proteins (Fig. 1A). Similarly, Ankyrons generated against the Delta variant, have binding scores for Delta spike protein that are significantly higher than those for the negative control (FcγR protein). In addition, Ankyrons, DT1 & DT3 cross react with the wild type and Omicron spike proteins (Fig. 1B). Finally, six of the 9 Ankyrons generated against the Omicron variant, have binding scores for Omicron spike protein that are significantly higher than those for the negative control (FcγR protein). Moreover, Ankyrons, Omc3 cross-reacts with the wild type spike protein while Omc9 cross-reacts with both wild type and Delta spike proteins (Fig. 1C).

**Fig 1.**
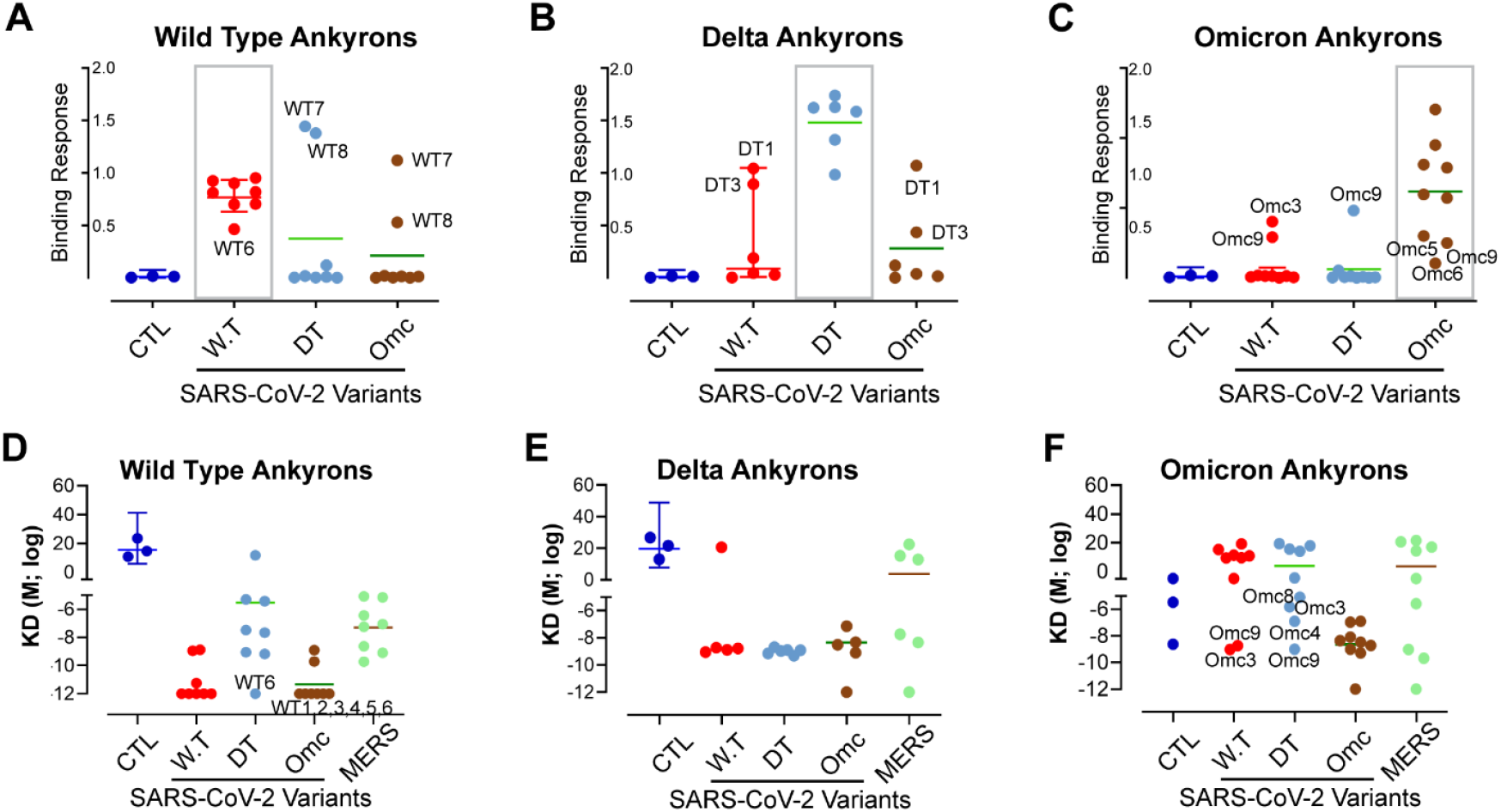
Identification of Ankyrons that directly interact with RBD protein of SARS-CoV-2 using the BLI binding kinetics assay. Binding Response based BLI assay was employed to monitor binding rate and specificity of three different Ankyrons: wild type (A), Delta (B) and Omicron (C) with immobilized spike S1+S2 trimer ECD proteins. These proteins included the wild-type strain (2019-nCoV) and two variants, Delta (B.1.617.2) and Omicron (B.1.1.529). Additionally, the binding kinetics between the S1+S2 variants (WT, Delta, and Omicron) and Ankyrons from wild type (D), Delta (E), and Omicron (F) were analyzed based on their affinity, represented by the KD value (equilibrium dissociation constant; M, mol/L) in an BLI analysis. The results also include a negative control targeting FcγR (CTL; blue), wild type (WT; red), Delta (DT; light blue), Omicron (Omc; brown), and MERS-CoV full-length trimer spike variant (MERS; green). Significance of differences between Ankyrons and trimeric spike proteins with respect to Ka, Kd, or KD was conducted using two-sided t test and adjusted for multiple comparisons using the Benjamini, Hochberg method with a false positive cutoff value of 0.10.

The KD values for Ankyrons generated against wild type, Delta, and Omicron spike proteins are shown in Fig. 1D-F. Compared to the negative control, all wild type Ankyrons bound with high affinity (low KD values) to wild type spike protein. In addition, the WT6 and WT1,2,3,4,5,6 exhibit cross-reactivity to Delta and Omicron spike proteins respectively (Fig. 1D). Similarly, all Delta Ankyrons bound with high affinity to the Delta spike proteins. Interestingly, most Delta Ankyrons also bound with high affinity to wild type and Omicron variants (Fig. 1E). The Omicron Ankyrons bound with high affinity to the Omicron spike proteins. The Omc3,9 and Omc3,4,8,9 exhibit cross-reactivity to wild type and Delta proteins respectively (Fig. 1F).

Although the KD and binding responses are consistent with respect to interactions between Ankyron and the spike protein variant used to generate the Ankyron, both parameters do not identify the same cross-reacting Ankyrons. We therefore used an analytical tool that identifies positive Ankyron-spike protein interactions based on conservative thresholds set for both binding response and KD (Fig. 2A-C). Ankyron-spike protein combinations that exceeded the threshold for both parameters were deemed positive. Based on this analysis we identified the following cross-reactive Ankyrons: (i) Wild type Ankyrons WT7 and WT8 bind to Delta and Omicron spike proteins. (ii) Delta Ankyrons DT1 and DT3 bind to wild type spike proteins while DT1 binds to both wild type and omicron spike proteins. (iii) No Omicron Ankyron bound to wild type spike protein, however, Omc9 bound with high affinity to the Delta spike protein.

**Fig 2:**
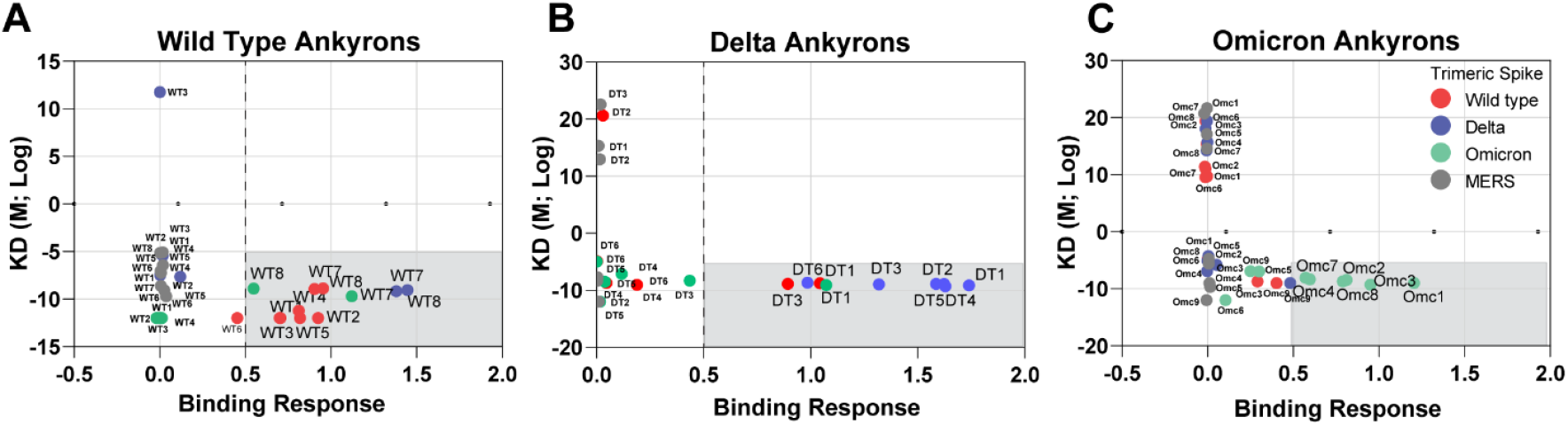
Approach for determining Ankyron-spike protein interaction based on binding response and KD value. We utilized an analytical tool to identify positive Ankyron-spike protein interactions, applying conservative thresholds for both binding response and KD values for Ankyron-spike protein interactions. Ankyrons which were generated against wild type (A), Delta (B) and Omicron (C) spike proteins were evaluated. Ankyron-spike protein combinations that exceeded the thresholds for both parameters were classified as target specific. Ankyrons that meet the criterion of KD < -5 (M; Log) and binding response > 0.5 are depicted in the gray box. The spike proteins used in the assay are: wild type (WT; red), Delta (DT; light blue), Omicron (Omc; brown), and MERS-CoV (MERS; green).

### Detection of specific and cross neutralizing activity of Ankyrons against SARS-CoV-2 infection

In addition to binding affinities, we determined the potential of Ankyrons to neutralize SARS-CoV-2 infection. The assessment was carried out in hACE2/TMPRSS2 HEK293 cells using a pseudovirus neutralizing assay expressing the wild type, Delta, and Omicron variants. As controls we used two monoclonal antibodies: (i) AcroB which has been demonstrated to neutralize wild-type, delta, and omicron SARS-CoV-2 infections^27^. (ii) C68 (Bamlanivimab) ^28, 29^ which is a potent inhibitor of wild type and delta but not the omicron SARS-CoV-2^29^. Neutralization of pseudovirus variants following pre-inoculation with Ankyrons that were generated against WT (WT1 to WT8), Delta (DT1 to DT6) and Omicron (Omc1 to Omc9) was evaluated using ACE2/TMPRSS2 overexpressing HEK293 cells. Neutralization of the wild type, delta, and omicron pseudoviruses is shown in Fig. 3A, B and C respectively. The IC_50_ values for each Ankyron-pseudovirus combination are reported in Table 1. A heat map (Fig. 3D) permits easy visualization of neutralization of viral infection by each of the Ankyrons. This heat map shows that Ankyrons neutralize the variant they were generated against with high potency (IC_50_, 0.56 to 16nM). In addition, Ankyrons WT7, WT8, DT1, DT3, and Omc9 neutralize SARS-CoV-2 infection of all variants.

**Fig 3.**
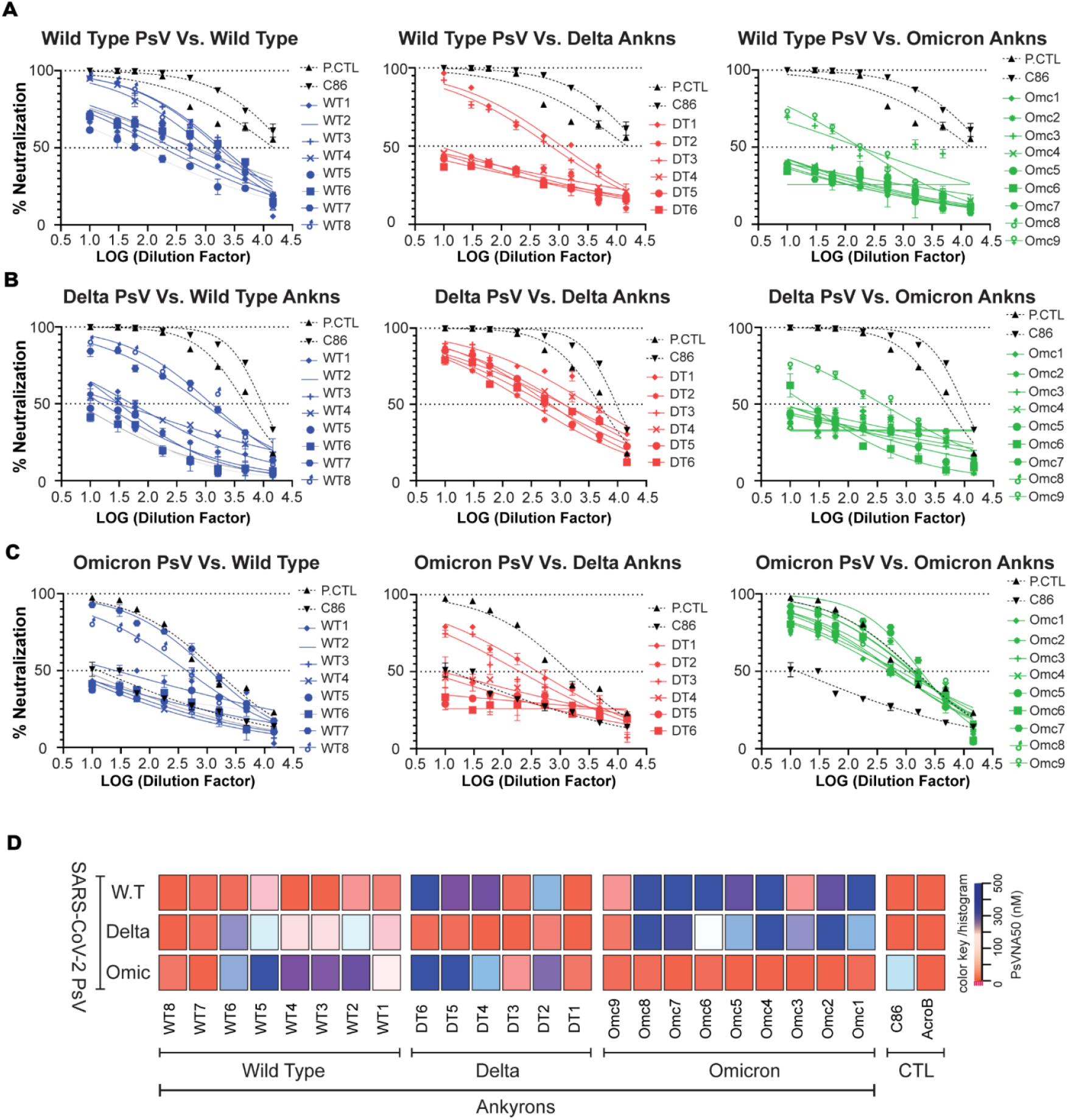
Ankyron-mediated neutralization of SARS-CoV-2 Pseudovirus variants. SARS-CoV-2 neutralizing titers for each Ankyron were determined using the pseudovirion neutralization assay in ACE2/TMPRSS2 over expressing HEK293 cells are depicted in panels A-C. The monoclonal antibodies ArcoB (black triangle) and C86 (inverted black triangle) are used as positive controls. (A) Wild type Pseudovirus (1.0 × 10^6^ RLU/mL) were used to infect ACE2/TMPRSS2 overexpressing HEK293 cells in presence of a 3-fold serial dilution of wild type (n=8), delta (n=6) and omicron (n=9) Ankyrons. (B) Delta pseudovirus (1.0 × 10^6^ RLU/mL) and (C) Omicron Pseudovirus (1.0 × 10^6^ RLU/mL) were also incubated with 3-fold dilutions of variant specific Ankyrons. The neutralization experiments were performed in triplicate and were repeated twice with similar results. (D) Heatmap showing the neutralization potency of specific Ankyrons against SARS-CoV-2 wild type, Delta, and Omicron variants. The IC_50_ value for each pseudovirus-Ankyron combination is shown, with pink, white, and dark-blue shading indicating high, intermediate, low, or no discernable binding, respectively.

### Ankyron-spike protein binding affinity and neutralization of SARS-CoV-2 variants

Viral neutralization depicted in Fig. 3 represents the clinically relevant effect of Ankyrons. However, BLI provides a high throughput assay to screen large numbers of Ankyrons. Such a high-throughput screening assay is required during the rapid selection of Ankyrons for neutralization of emerging variants. We find that the Ankyrons that neutralize the SARS-CoV-2 virus bind to the variant that was neutralized with high affinity (KD value in Table 1). Thus, rapid high throughput screening of Ankyrons is an appropriate strategy for a tiered approach (i.e., including only those Ankyrons in a neutralization assay that bind with high affinity based on a pre-determined cut-point).

## Discussion

Prophylactic vaccines continue to be an effective and cost-effective public health response to pandemics such as COVID-19. Nonetheless, additional clinical tools are needed because: (i) An epidemic/pandemic due to a new virus lacks a vaccine and millions of patients need to be treated. (ii) A vaccine may not be universally effective. (iii) As new viral strains emerge, vaccines developed against previous strains may not be effective. (iv) Vaccine hesitancy leads to a group that is not vaccinated and may need alternate clinical interventions. Recombinant mAbs offer one such alternative but have limitations (see Introduction). Here, we present an alternative approach that uses synthetic molecules, called Ankyrons, that can be generated to bind with high affinity to diverse targets akin to antibodies. We evaluate Ankyrons that were generated against various strains of the spike protein of SARS-CoV-2.

Ankyrons were generated against the following variants of the spike protein of SARS-CoV-2: Wild type (WT1-WT8), (iii) Delta (DT1-DT6), and Omicron (Omc1-Omc9). An Ankyron that targeted the FcγR was used as a negative control. Using BLI we demonstrated that the Ankyron generated against FcγR did not bind SARS-CoV-2 spike proteins. Similarly, none of the Ankyrons generated against the SARS-CoV-2 spike proteins bound to the spike protein from the closely related coronavirus, MERS. All Ankyrons bound with high affinity to the spike protein variant used to generate the Ankyron. Together these data show that the Ankyrons described in this study are specific to the SARS coronavirus.

A consistent challenge in preventing and treating COVID 19 has been the periodic emergence of new variants^30^. Biologics such as monoclonal antibodies that are effective against one strain of SARS-CoV-2 are often not effective against another variant ^31-33^. A potential advantage of using Ankyrons as antiviral agents is that numerous unique Ankyrons are simultaneously generated against a single target. It is thus likely that some of these Ankyrons could target multiple spike protein variants. We evaluated the cross reactivity of the 23 anti-SARS-CoV-2 spike protein Ankyrons that were generated in this study. We adopted a conservative approach wherein an Ankyron was considered cross reactive if pre-determined thresholds were met for two parameters, binding response, and KD. The BLI method provides multiple parameters and binding response, and KD are both used to determine interactions between a ligand and an analyte ^25, 26^. The Ankyrons WT7, WT8, DT1, DT3, Omc3 and Omc9 were cross-reactive with all three strains of SARS-CoV-2 spike proteins tested, i.e., wild type, Delta, and Omicron.

Binding affinities carried out in a high throughput platform like BLI provide a useful means of screening Ankyrons. Binding of the Ankyron is a necessary condition for viral neutralization but not sufficient. Thus, the results of the binding assay do not establish whether the bound Ankyrons will neutralize SARS-CoV-2 infections. We carried out a neutralization assay using pseudoviruses of all three SARS-CoV-2 strains. Using this assay, we demonstrate that Ankyrons WT7, WT8, DT1, DT3 and Omc9 exhibit cross-reactivity and can neutralize all three strains of SARS-CoV-2.

This study demonstrates that emerging molecular entities called Ankyrons can be leveraged to rapidly develop neutralizing agents for viruses that cause epidemics and pandemics. Ankyrons were obtained in high throughput *in vitro* ribosome display against different strains of the SARS-CoV-2 spike proteins. Within this pool were multiple Ankyrons that could neutralize all three strains of SARS-CoV-2 used in this study. The potency of cross-reactive Ankyrons is high (IC50, 0.089 to 5.25nM), which proves the principle that this Ankyron technology could produce cross-reactive binders rapidly. The technology permits additional rounds of design and evaluation of multi-valent moieties that can either increase the potency and/or expand the repertoire of SARS-CoV-2 strains that can be neutralized. Furthermore, it is important to emphasize that we have evaluated only a small fraction of the total number of anti-spike protein Ankyrons that were generated. We found that neutralizing Ankyrons were a subset of Ankyrons that were determined to be binders based on the conservative criteria for identifying binding-Ankyrons. Thus, strategies that use an automated high-throughput method identifying binding-Ankyrons can be incorporated into a tiered workflow that can screen very large numbers of Ankyrons. The Ankyrons that meet the signal-to-noise threshold for binding to a specific target by immunoassay can then be used in the more labor and cost-intensive neutralizing assay. By screening Ankyrons obtained through ribosome display for binding against different variant targets it is possible to evaluate cross reactivity quickly. The process of generating new target binding Ankyrons can typically be accomplished withing 6-8 weeks, including expression and purification of several engineered variants for further testing, following the availability of the targets themselves.

Taken together, our study describes an efficient workflow for the generation and evaluation of Ankyrons against a viral target. Using the example of SARS-CoV-2, we demonstrate that multiple Ankyrons can be raised against each strain of the spike protein and that some of these Ankyrons can cross-react to and neutralize additional strains of the SARS-CoV-2 virus. Ankyrons provide a feasible, rapid, and cost-effective solution to developing drugs that can combat existing and emerging strains of the SARS-CoV-2 virus. Given the complexity of the SARS-CoV-2 spike proteins this raises the hope that Ankyrons could also be useful for rapidly developing new research tools for studying emerging infectious diseases, including for tens and hundreds of different targets in parallel, if necessary, with the optional potential for developing them further into diagnostic and even therapeutic applications, such as in passive immunization.

## Material and methods

### Reagent and supplies

Phosphate buffered saline (PBS) tablets (Sigma P4417), Tween-20 (Fisher BP337), Trypsin 0.25% EDTA (Fisher 25200056), Luciferase Assay system (Promega E1501), Luciferase Passive Lysis buffer (Promega E1941), SARS-CoV-2 spike (WT) Fluc-GFP pseudovirus (ACROBiosystems PSSW-HLGB001), SARS-CoV-2 spike (Delta) Fluc-GFP Pseudovirus (ACROBiosystems PSSD-HLGB002), SARS-CoV-2 spike (Omicron) Fluc-GFP Pseudovirus (ACROBiosystems PSSO-HLGB003), Anti-SARS-CoV-2 spike RBD Neutralizing Antibody, Chimeric mAb, Human IgG1 (ACROBiosystems S1N-M122), 96-well ELISA plate (Fisher C37000), Bovine Serum Albumin (Akron Biotech AK1391), SARS-CoV-2 (2019-nCoV) Spike S1+S2 ECD (R683A, R685A,F817P,A892P,A899P,A942P,K986P,V987P)-His (SinoBiological 40589-V08H4), SARS-CoV-2 B.1.617.2 Spike S1+S2 trimer protein ECD-His tag (SinoBiological 40589-V08H10), SARS-CoV-2 B.1.1.529 (Omicron) S1+S2 trimer protein ECD-His tag (SinoBiological 40589-V08H26), Human ACE2 293T Cell line (Takara Bio 631289), Octet Anti-Penta-His (HIS1K) sensor tips (Sartorius ForteBio 18-5120), Octet Streptavidin (SA) sensor tips (Sartorius ForteBio 18-5019), tilted bottom (TW384) microplates (Sartorius ForteBio 18-5080), DMEM+GlutaMAX (Gibco 10569-010), Heat inactivated Fetal Bovine Serum (Gibco 16140-071).

### Generation of Ankyrons

Ankyrons specific to the relevant SARS-CoV-2 spike protein variants were obtained from ProImmune Ltd (Oxford. UK). According to the company, Ankyrons are generated by using the ProImmune Ankyron Teralibrary in *in vitro* ribosome display which was carried out with each of the three SARS-CoV-2 spike protein variants in turn to generate a number of Ankyron clones with the ability to bind their cognate target. A final selection of unique Ankyron clones was isolated, based on their performance and specificity in target-binding immunoassays. Selected Ankyrons were expressed with a histidine tag as well as either a V5 epitope tag or a tag to allow for site-specific biotinylation, where required. The cloned Ankyrons were then overexpressed and purified using standard chromatography techniques. Overall the following Ankyrons specific to variants of the spike protein of SARS-CoV-2 were provided with the signal to noise (SNR) ratio for each clone determined from the initial immunoassay also shown in italics: wild type [8 Ankyrons: WT1 (clone AZ41031; SNR*1108*), WT2 (clone AZ40748; SNR*941*), WT3 (clone AZ41032; SNR*875*), WT4 (clone AZ41033; SNR*730*), WT5 (clone AZ40754; SNR*491*), WT6 (clone AZ41035; SNR*450*), WT7 (clone AZ40758; SNR*359*), WT8 (clone AZ41041; SNR*307*)], delta [6 Ankyrons: DT1 (clone AZ40860; SNR*1744*), DT2 (clone AZ40861; SNR*768*), DT3 (clone AZ40862; SNR*570*), DT4 (clone AZ40864; SNR*433*), DT5 (clone AZ40880; SNR*203*), DT6 (clone AZ40883; SNR*163*)], and omicron [9 Ankyrons: Omc1 (clone AZ40218; SNR*7355*), Omc2 (clone AZ41007; SNR*5241*), Omc3 (clone AZ41011; SNR*4183*), Omc4 (clone AZ41013; SNR*4094*), Omc5 (clone AZ41014; SNR*3916*), Omc6 (clone AZ41015; SNR*3266*), Omc6 (clone AZ40220; SNR*3080*), Omc7 (clone AZ41018; SNR*2593*), Omc8 (clone AZ40225; SNR*2324*)]. In addition, for use as negative controls Ankyrons that target the Fcγ receptor (FcγR) FCR1 (clone AD40138; SNR*1412*), FCR2 (clone AD40145; SNR*114*), FCR3 (clone AT30369; SNR*476*) were also provided.

### BioLayer interferometry assay (BLI)

Bio-Layer Interferometry (BLI), a label-free technology, was used for measuring the interactions of SARS-CoV-2 spike trimer proteins with each Ankyron peptides. The affinity measurements were performed with Octet RED96 instrument with streptavidin (SA) biosensor tips (ForteBio, Inc., Menlo Park, CA, USA) and Octet System Data Acquisition software version 8.2 (ForteBio; https://www.fortebio.com/products/octetsystems-software). Biotinylated Ankyrons (ProImmune) were loaded onto streptavidin biosensor at 25nM concentration, recombinant wild-type SARS-CoV-2 full length spike trimer protein, Delta full length trimer spike variant, Omicron B.1.1.529 full length trimer spike variant and MERS full length trimer spike variant (ACROBiosystems Cat No. SPN-M52H4) were loaded at various concentration (50nM to 0.625nM by serial dilution) for real time association and dissociation analysis using the ForteBio Octet RED96 system. BLI assay buffer consists of 0.1% BSA and 0.02% Tween 20 in PBS, which was 0.22-um filtered. Before use, anti-biotin (streptavidin, SA) biosensors were loaded into the columns of a biosensor holding plate and pre-hydrated in BLI assay buffer for 10min. Tilted bottom 96-well plates were loaded with 200uL per well. The assay plate was prepared as follows: column 1 (BLI assay buffer), Column 2-5 (each 25nM Ankyrons in BLI assay buffer), Column 8 (BLI assay buffer for washing 1), Column 9 (serial diluted SARS-CoV-2 trimeric RBD from 50nM to 0.78nM), Column 10 (BLI assay buffer for washing 2). The BLI-ISA method was set as follows. Baseline1 (60 s) in column 1 (Equilibration), Loading (150 s or 600 s) in column 2-5 (each different Ankyrons, respectively), Baseline2 (60 s) in column 8 (Wash), Association (500 s) in column 9 (Trimeric SARS-CoV-2 Spike proteins), Dissociation (700 s) in column 10. The assays were maintained at a temperature of and the speed of 30°C and 1000 rpm respectively. The typical resulting capture levels of eight biosensors were in the range of 0.4–0.5 nm. The association (kon) and dissociation (kdis) were then established by dipping the biosensors for 10 mins in various concentrations of Ankyron samples dispensed in 96-microwell plates. Data were processed and analyzed using the Octet data analysis software version 7.0 (ForteBio). The binding profile of each Ankyron was shown as “nm” shift. This shift is a comparison of the shift in the interference patterns of light reflected from a reference layer within the biosensor versus molecules as the bind to the biosensor tip. The results were summarized as KD which was calculated from “KD□=□kon/kdis”, where kon is the ‘on rate’ or association and kdis is the ‘off rate’ or dissociation.

### SARS-CoV-2 Pseudovirus Production and Neutralization Assay

A pseudovirus neutralization assay was performed to detect the neutralizing activity of all Ankyrons that were specifically targeting against SARS-CoV-2 infection, or their cross-neutralizing activity against SARS-CoV-2 infection. ACROBiosystems antibody (Anti-SARS-CoV-2 Spike RBD Neutralizing Antibody, Chimeric mAb, Human IgG1), C86 (Ly-CoV555; homemade Bamlanivimab) and a MERS-CoV RBD-specific mAb were used as controls (Tai et al., 2020; Du et al., 2016). SARS-CoV-2 Fluc-GFP Pseudovirus for Wild Type, Delta and Omicron uses pseudotyped HIV-1 virus with firefly luciferase and green fluorescent protein (GFP) gene as the backbone and takes SARS-CoV-2 spike protein as its envelope protein (ACROBiosystems PSSW-HLGB001: WT, PSSD-HLGB002: Delta, PSSO-HLGB003: Omicron). Neutralization assays were performed on 293T cells transiently transfected or transduced with ACE2 and TMPRSS2 genes for stable expression. The Ankyron dilution was tripled; 1:20 (100nM), 1:60 (33.3nM), 1:180 (11.1nM), 1:540 (3.7nM), 1:1620 (1.24nM), 1:4860 (0.41nM), 1:14580 (0.137nM), 1:43740 (0.045nM). Briefly, pseudoviruses with titers of approximately 10^6^ RLU/mL of luciferase activity were incubated with triple diluted Ankyrons for one hours at 37°C. Pseudovirus and Ankyrons or antibody mixtures (100ul) were then inoculated onto 96well plates that were seeded with 2.5 × 10^5^ cells/well on day prior to infection. Pseudovirus infectivity was scored 48h later for luciferase activity. The Ankyrons or antibody concentration causing a 50% reduction of RLU compared to control (ID50 or IC50, respectively) were reported as the neutralizing antibody titers. Titers were calculated using a nonlinear regression curve fit (GraphPad Prism software Inc., La Jolla, CA). The mean 50% reduction of RLU compared to control from at least two independent experiments was reported as the final titer. Treated cells were lysed in cell lysis buffer (Promega, Madison, WI) 72 h after infection. Luciferase substrate (Promega) was added to the lysed cell supernatants, which were detected for relative luciferase activity using the (BioTek Synergy HTX Multimode Reader, Agilent, Santa Clara, CA). The percent (%) pseudovirus neutralization was calculated.

### Statistical analyses

Biomolecular binding kinetics parameters (association rate constant (ka) [1/Ms]; dissociation rate constant (kd) [1/s]; affinity rate constant (KD) [M]) were analyzed using Octet System Data Analysis software version 8.2 (Pall ForteBio; https://www.fortebio.com/products/octet-systems-sofware). Only data sets with curve ft R2 values greater than 0.95 were included in further analyses as per the manufacturer’s recommendation. Significance of differences between Ankyrons and trimeric spike proteins with respect to Ka, Kd, or KD was conducted using two-sided t test and adjusted for multiple comparisons using the Benjamini, Hochberg method with a false positive cutoff value of 0.10. Testing the significance of fold change of KD comparing controls (AcroB or C38) was achieved through OneWay ANOVA and controlled for variations using multiple regressions.

## Acknowledgments

We thank Dr. Daniel Lagasse for discussions and suggestions.

## Funding

This work was supported by intramural research funding from the US Food and Drug Administration.

## Declaration of Competing Interest

Nicholas C Hurst is an employee of ProImmune Ltd. Jeremy W Fry is an employee and shareholder of ProImmune Ltd. Nikolai F Schwabe and Linda C C Tan are founders and shareholders of ProImmune Ltd.

## Notes

### Summary of Updates

Textual changes have been made to enhance the clarity of the manuscript.

