## Supplementary information for "Functional Activity and Binding Specificity of small Ankyrin Repeat Proteins called Ankyrons against SARS-CoV-2 variants"

**Supplementary Figure. 1.**

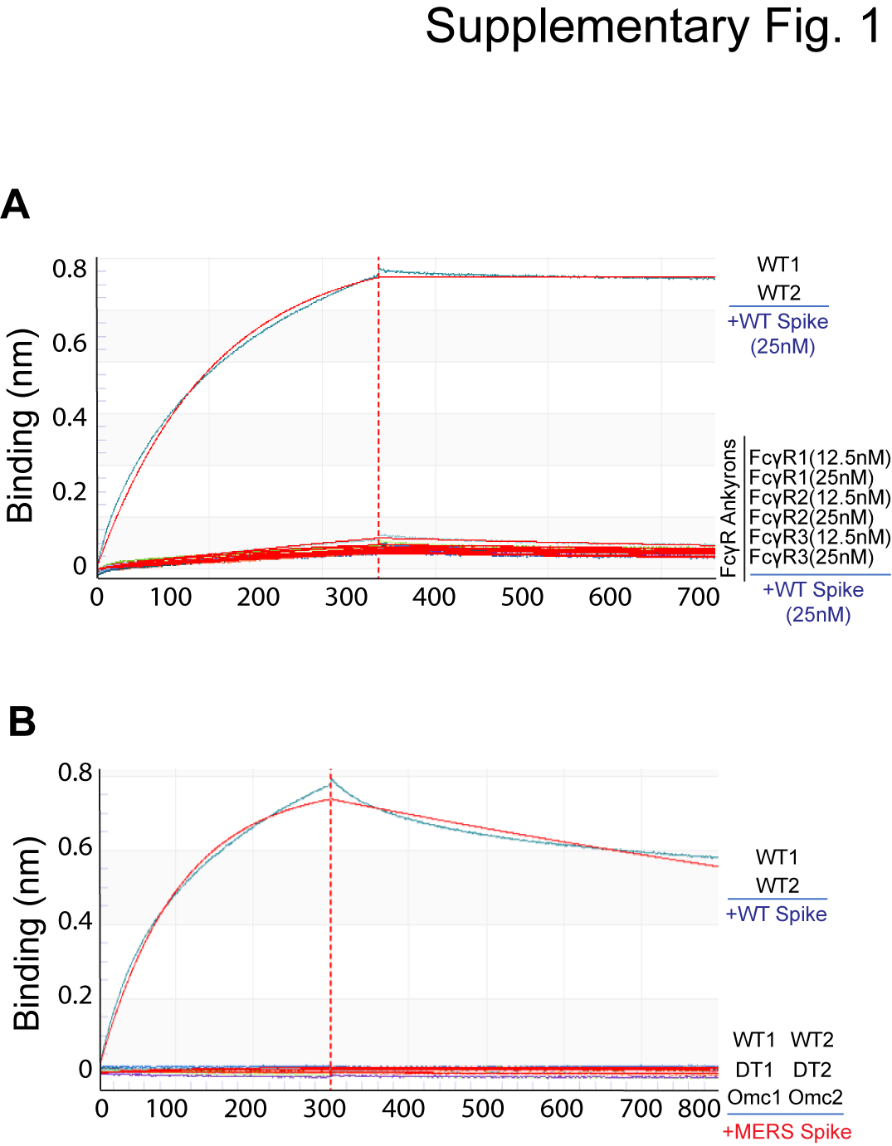

**Supplementary Figure. 1. Verification of binding specificity and cross-reactivity of Ankyrons with SARS-CoV-2 spike protein.**

To measure the binding affinity, specificity, and cross-reactivity of Ankyrons with SARS-CoV-2 variants, two individual control experiments were performed for the control groups. Three different types of Ankyrons (FcγR1, FcγR2 and FcγR3) against for Fcγ receptor were tested at two different concentrations (12.5nM and 25nM) with the S1+S2 trimeric protein wild-type (A) and two variants [Delta (B.1.617.2) and Omicron (B.1.1.529)] as well (data not shown); Additionally, two wild type Ankyrons (WT1 and WT2), selected for highest binding affinity, were used as positive controls for this experiment. (B) 25nM of Target specific Ankyrons (WT1, WT2, DT1, DT2, Omc1 and Omc2) were tested against 25nM of the MERS spike protein using a BLI assay, and the WT1 and WT2 Ankyrons were also used as positive controls in this BLI experiment alongside the wild-type trimeric spike.

**Supplementary Figure. 2**.

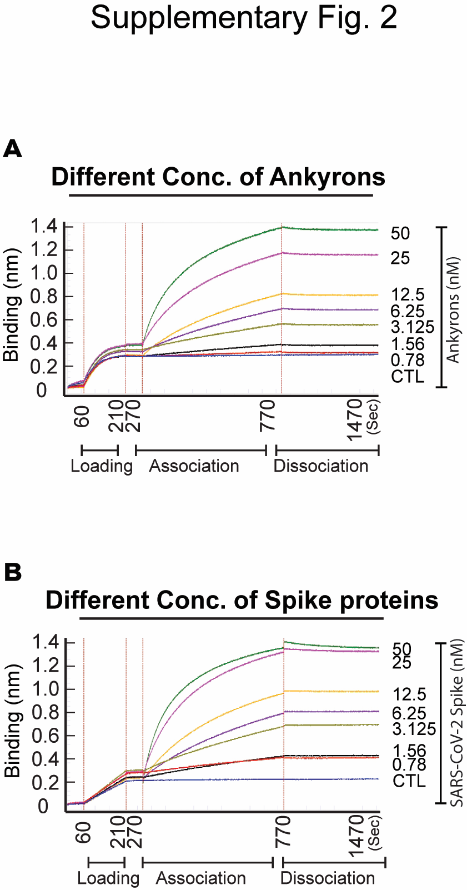

**Supplementary Figure. 2. Optimization of BLI analysis using various concentration of analytes (Ankyrons) and Ligands (trimeric spike proteins variant)**

To refine the BLI analysis conditions, two distinct experimental setups were evaluated. (A) Spike trimer was loaded onto biosensor tips at a constant concentration of 25nM, and the association phase was monitored as Ankyrons, serially diluted from 50nM to 0.78nM, were introduced. (B) In a reverse setup, Ankyrons were fixed at 25nM and bound to the biosensor, with the association and dissociation phases tracked as the Spike trimer, serially diluted from 50nM to 0.78nM, was added. FcrR-specific Ankyrons served as the control. The association phase lasted 500 second, followed by a 700 second dissociation phase. Binding was measured as a wavelength shift (Δλ; in nanometers [nm]) detected by the Octet instrument.

**Supplementary Figure. 3.**

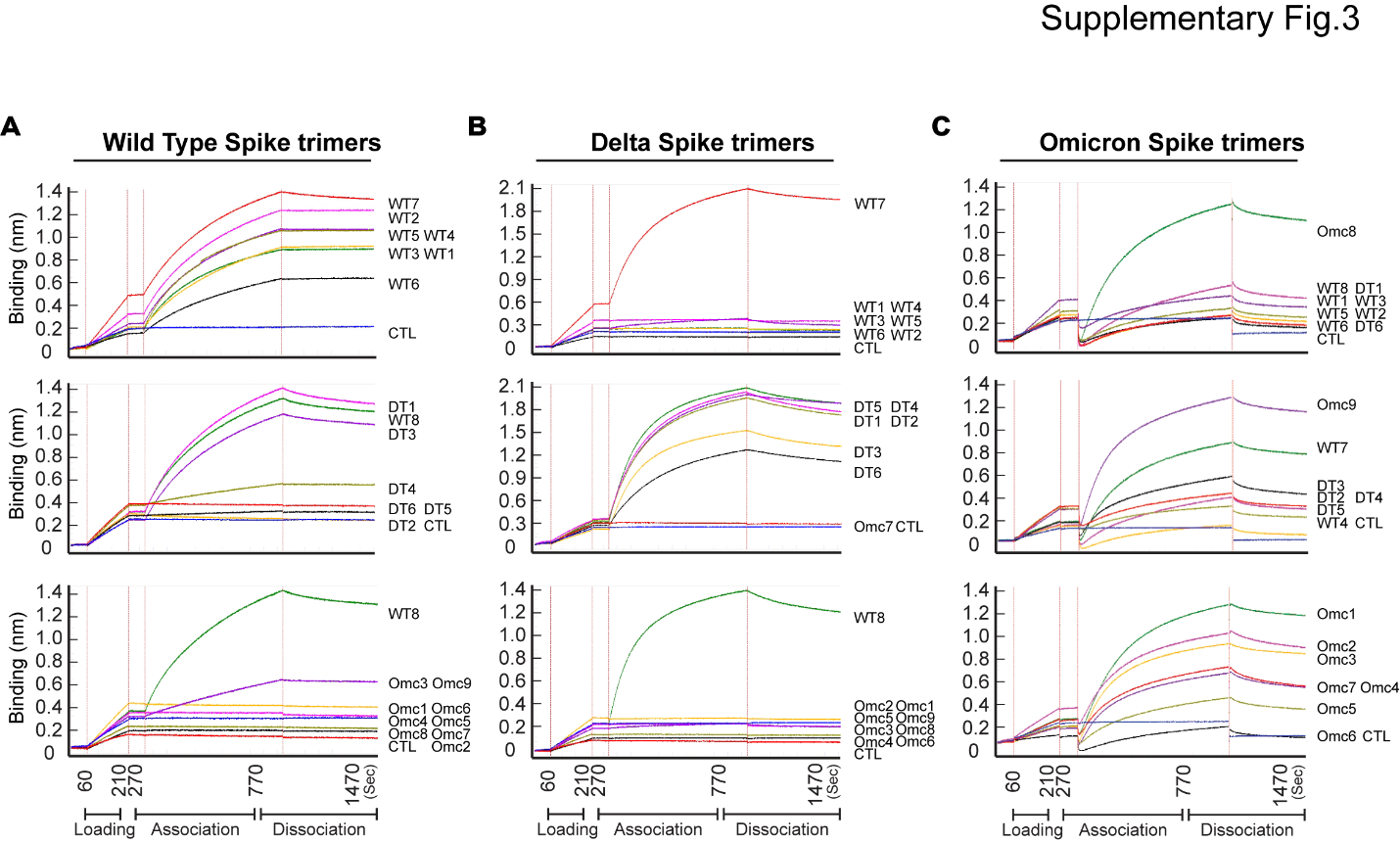

**Supplementary Figure. 3. Biolayer interferometry (BLI) analysis for identifying specific Ankyrons that directly interact with RBD protein by BLI binding kinetics assay.**

To confirm the target-specific binding of each Ankyrons, wild type (n=8), Delta (n=6), and Omicron (n=9) Ankyrons were analyzed for their binding affinity and specificity to the S1+S2 trimeric full-length spike RBD proteins of three SARS-CoV-2 variants [wild type, Delta (B.1.617.2), and Omicron (B.1.1.529)] using BLI analysis. (A) Wild type trimeric spike proteins at a concentration of 25nM were initially loaded onto the biosensor, followed by testing the binding affinity of all Ankyrons at the same 25nM concentration. (B) The Delta (B.1.617.2) spike trimer and (C) the Omicron (B.1.1.529) spike trimer was similarly tested at a 25nM dilution for binding with all specific Ankyrons. This analysis revealed specific binding as well as nonspecific cross-reactivity between certain Ankyrons and the target spike variants. The association phase lasted 500 seconds, followed by a 700-second dissociation phase. Binding was measured as a wavelength shift (Δλ; in nanometers [nm]) detected by the Octet instrument.

**Supplementary Table. 1**.

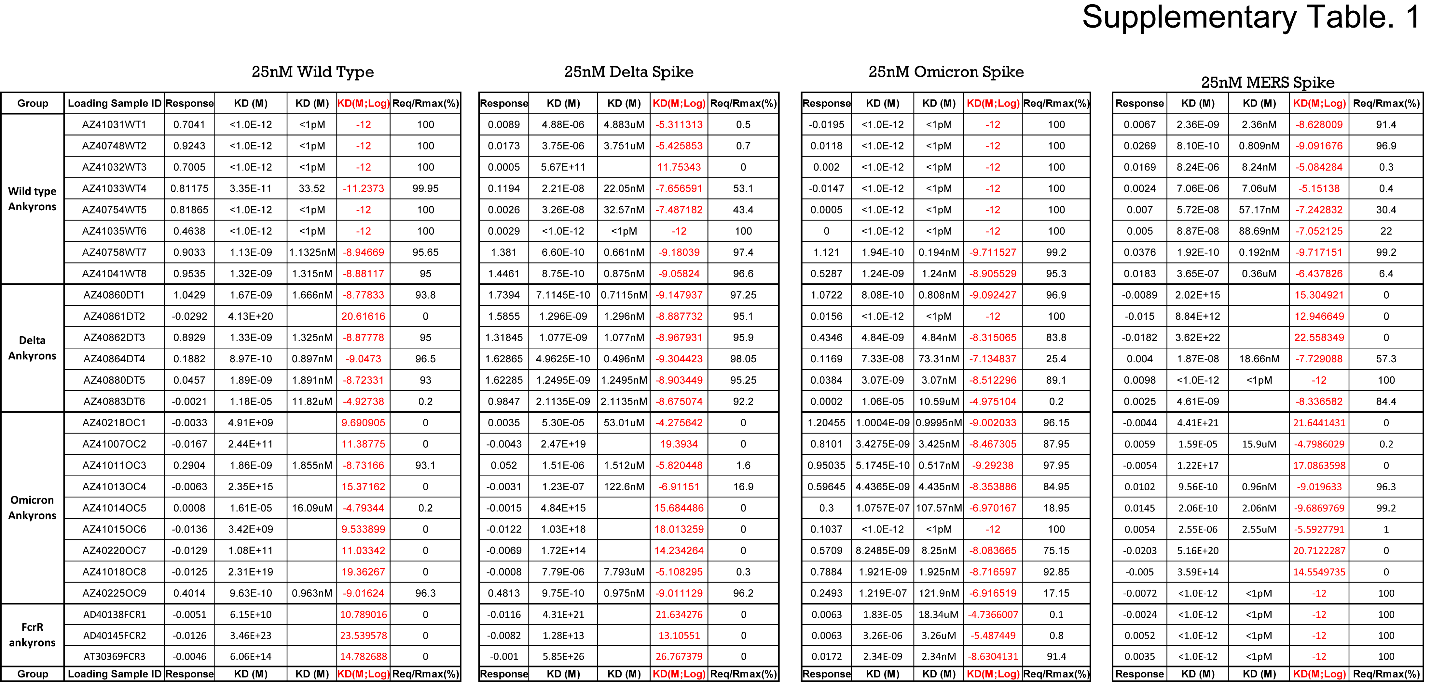

**Supplementary Table. 1. Summary of Biolayer interferometry data for binding affinity and specificity between Ankyrons and SARS-CoV-2 spike variants.**

KD (equilibrium affinity constant, M, mol/L; range: 1mM – 10pM); equilibrium binding signal (R_eq_); and maximum binding signal (R_max_). The ratio of R_eq_ to R_max_ depends on the analyte concentration relative to the KD. A mean KD of <1E-12 M was not calculated for variants due to the strong binding between analytes and ligands.
